# Peri-saccadic orientation identification performance and visual neural sensitivity are higher in the upper visual field

**DOI:** 10.1101/2022.07.05.498850

**Authors:** Alessio Fracasso, Antimo Buonocore, Ziad M. Hafed

## Abstract

Visual neural processing is distributed among a multitude of sensory and sensory-motor brain areas exhibiting varying degrees of functional specializations and spatial representational anisotropies. Such diversity raises the question of how perceptual performance is determined, at any one moment in time, during natural active visual behavior. Here, exploiting a known dichotomy between the primary visual cortex and superior colliculus in representing either the upper or lower visual fields, we asked whether peri-saccadic orientation identification performance is dominated by one or the other spatial anisotropy. Humans (48 participants, 29 females) reported the orientation of peri-saccadic upper visual field stimuli significantly better than lower visual field stimuli, unlike their performance during steady-state gaze fixation, and contrary to expected perceptual superiority in the lower visual field in the absence of saccades. Consistent with this, peri-saccadic superior colliculus visual neural responses in two male rhesus macaque monkeys were also significantly stronger in the upper visual field than in the lower visual field. Thus, peri-saccadic orientation identification performance is more in line with oculomotor, rather than visual, map spatial anisotropies.

**Significance statement:** Different brain areas respond to visual stimulation, but they differ in the degrees of functional specializations and spatial anisotropies that they exhibit. For example, the superior colliculus both responds to visual stimulation, like the primary visual cortex, and controls oculomotor behavior. Compared to the primary visual cortex, the superior colliculus exhibits an opposite pattern of upper/lower visual field anisotropy, being more sensitive to the upper visual field. Here, we show that human peri-saccadic orientation identification performance is better in the upper compared to the lower visual field. Consistent with this, monkey superior colliculus visual neural responses to peri-saccadic stimuli follow a similar pattern. Our results indicate that peri-saccadic perceptual performance reflects oculomotor, rather than visual, map spatial anisotropies.

## Introduction

Natural active visual behavior is characterized by frequent saccadic eye movements used to scan our environment. At the time of saccades, vision is not necessarily completely halted (Ross et al., 2001; De Pisapia et al., 2010; Fracasso et al., 2015; Binda and Morrone, 2018; Grujic et al., 2018; Zimmermann, 2020), but it is certainly altered. For example, visual sensitivity can be strongly suppressed peri-saccadically (Latour, 1962; Matin, 1974; Diamond et al., 2000; Idrees et al., 2020a), and spatial localization perceptual performance is grossly distorted (Ross et al., 1997; Kaiser and Lappe, 2004; Binda et al., 2009). Temporal judgements are additionally affected by saccades (Morrone et al., 2005). This evidence suggests that peri-saccadic vision is phenomenologically fundamentally different from vision during steady-state gaze fixation (Wurtz and Goldberg, 1971, 1972; Bruce and Goldberg, 1985; Russo and Bruce, 1993, 2000; Stanton et al., 2005; Hafed and Krauzlis, 2012).

The fact that peri-saccadic vision transpires at the same time as saccade motor commands leads to a question about the neural substrates supporting this special, albeit fleeting, kind of vision. In particular, it is well known that perceptual performance during steady-state gaze fixation is generally superior in the lower visual field (Talgar and Carrasco, 2002; Barbot et al., 2021), and increasing evidence suggests that the primary visual cortex (V1) exhibits neural tissue anisotropies that might explain such perceptual asymmetry (Benson et al., 2021; Kupers et al., 2022). On the other hand, the superior colliculus (SC) visual representation preferentially favors the upper visual field instead (Hafed and Chen, 2016), with neurons exhibiting higher and earlier visual sensitivity for stimuli above the retinotopic horizon than below it. If both V1 and SC neurons interact to coordinate visually-guided behavior, how might such divergent anisotropies in these two functionally and anatomically related brain areas determine perceptual performance, and particularly during the peri-saccadic interval? Answering this and related questions is important for better understanding how functional specializations in different visual and motor structures (Previc, 1990) can all work together to give rise to coherent behavioral outcomes.

We approached this problem by studying peri-saccadic orientation identification performance. It is generally accepted that the sensitivity of the visual system to brief peri-saccadic flashes is strongly suppressed (Latour, 1962; Diamond et al., 2000; Idrees et al., 2020b). However, residual visual processing still takes place at the time of saccades (De Pisapia et al., 2010; Fracasso et al., 2015; Fracasso and Melcher, 2016; Fabius et al., 2022), allowing us to ask whether such processing is more sensitive in the upper or lower visual fields. We first asked human subjects to generate horizontal saccades, and we presented upper or lower visual field peri-saccadic flashes, which were near the vertical retinotopic meridian at the time of peak saccadic suppression. We found that residual orientation identification performance was significantly higher in the upper visual field than in the lower visual field. This result was categorically different from our expectation that orientation identification performance should have been better, or at least the same, in the lower visual field (Kristjansson and Sigurdardottir, 2008; Barbot et al., 2021), an expectation that we also confirmed with our own control gaze fixation experiments in this study. However, this higher peri-saccadic orientation identification performance in the upper visual field was in line with the anisotropy that exists in the oculomotor system, symbolized by the SC’s preference for upper visual field stimuli (Hafed and Chen, 2016). Since it is still not known how peri-saccadic SC visual neuronal sensitivity operates (i.e. whether it still prefers the upper visual field), we next inspected SC visual responses in two rhesus macaque monkeys around the time of saccades. We found that SC peri-saccadic neuronal responses to visual stimuli were again still higher in the upper rather than the lower visual field. Our results suggest that peri-saccadic orientation identification performance reflects oculomotor, rather than visual, map anisotropies. This observation might imply prioritization for detecting extra-personal stimuli for rapid orienting or evasive responses exactly at the time at which perception may be most compromised by saccades.

## Materials and methods

Experiments 1 and 2, along with a control study associated with them, were psychophysical experiments on human participants. The third experiment consisted of analyzing neurophysiological recordings from two rhesus macaque monkeys. The human experiments (including the control) were approved by the University of Glasgow Research Ethics Committee, and the participants received compensation of £6 per testing hour. Written informed consent was also obtained, in accordance with the 1964 Declaration of Helsinki. The monkey experiments were approved by the Regierungspräsidium Tübingen, under licenses CIN3/13 and CIN04/19G, and the experiments complied with European and national laws governing animal research.

A total of 48 human subjects aged between 18 and 38 years took part in the human behavioral experiments (Experiment 1: 20 subjects, 14 females; Experiment 2: 14 subjects, 8 females; Control study: 14 subjects, 7 females). All subjects self-reported as being free from neurological impairments. All subjects also had normal or corrected to normal vision and were naïve to the purposes of the experiment. The neurophysiological analyses were performed on an existing data set from (Chen and Hafed, 2017), which we re-analyzed here from the perspective of visual field asymmetries. The monkeys in that study were two adult, male rhesus macaques aged 7 years.

In what follows, we first describe the human experiments, and we then report on the neurophysiological analyses.

### Experimental design and statistical analysis

In humans, for the main experiments (Experiment 1 and 2), visual stimuli were designed using two variables with two levels each: visual field location (upper or lower visual field with respect to the line of sight) and attentional allocation (diffuse or focused on the expected target location). In the control experiment, visual stimuli were varied along three variables: visual field location (upper or lower), eccentricity (3 degrees, deg, or 7 deg), and spatial frequency (0.9 cycles per degrees, cpd, or 6 cpd).

In monkeys, stimuli were presented in different positions within the visual field: upper or lower with respect to fixation.

All statistical analyses are described in detail at the relevant locations in the following methodological sub-sections. We provided descriptive statistics throughout the Results section, and all figures include measures of confidence across our recorded samples.

### Human laboratory setup and behavioral tasks

Stimuli were presented on a 24-inch LCD monitor (1920 x 1024 pixels) at 60 Hz. Subjects were seated with their head resting on a chin and forehead rest to reduce head movements. Eyes were horizontally and vertically aligned with the center of the screen at a distance of 65 cm. Eye movements were recorded with the EyeLink 1000 system (detection algorithm: pupil and corneal reflex; 1000 Hz sampling; saccade detection was based on 30 deg/s velocity and 9500 deg/s^2^ acceleration thresholds). Subjects’ responses were recorded on a standard keyboard. A five point-calibration on the horizontal and vertical axes was performed at the beginning of each experimental run. The programs for stimulus presentation and data collection were written in MATLAB (MathWorks) using the Psychophysics Toolbox Version 3 (Brainard and Vision, 1997; Pelli and Vision, 1997), and Eyelink Toolbox extensions (Cornelissen et al., 2002).

Stimuli included a fixation point measuring 0.7 deg, which was jumped to instruct saccade generation. Target stimuli were gabors with a spatial frequency of 0.9 cpd and a gaussian envelope with σ 3.5 deg (see Fig. 1A in Results). Distractor stimuli consisted of the sum of two gabors (one horizontal and one vertical), tilted by 45 degrees (see Fig. 1A in Results). In the control experiment requiring only gaze fixation, we generated gabor stimuli with a spatial frequency of 0.9 cpd and also 6 cpd, both having a gaussian envelope with σ 3.5.

**Figure 1.**
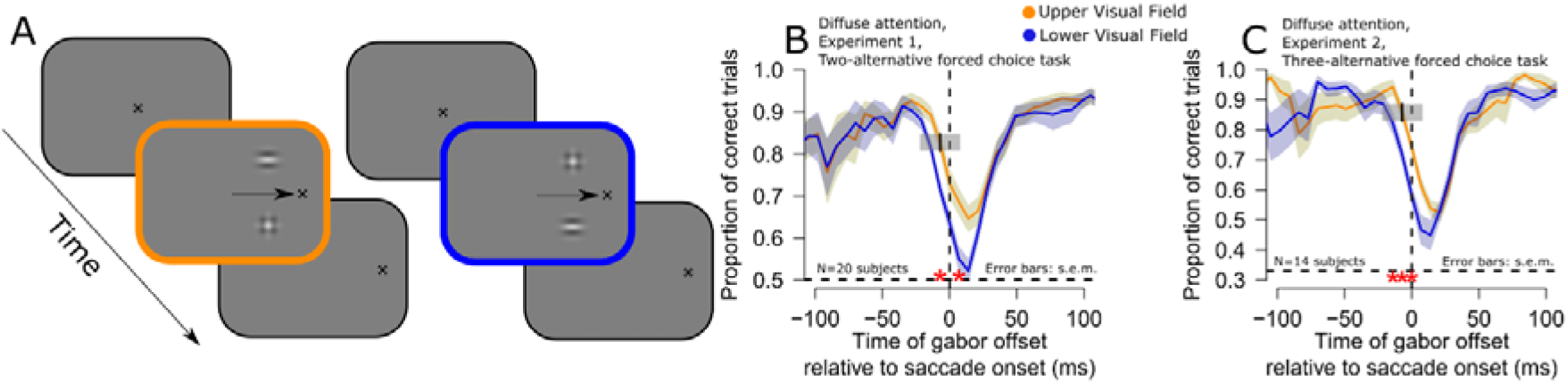
Better peri-saccadic orientation identification performance in the upper visual field. **(A)** Subjects generated ∼12 deg horizontal saccades (schematized by an arrow in the figure). At different times relative to saccade onset, two image patches appeared briefly, one above and one below the horizontal meridian (Materials and methods). One patch was an oriented gabor grating (the target), and the other was a distractor with no orientation information. The subjects reported the orientation of the target gabor, and we assessed whether the subjects’ responses were better when the target appeared in the upper (yellow) versus lower (blue) visual field. **(B)** Time course of orientation identification performance relative to saccade onset for targets in the upper (yellow) or lower (blue) visual field in Experiment 1 (diffuse attention condition; Materials and methods). Red asterisks indicate significant differences between the two curves (GLMM, main effect of target gabor grating location, p<0.01). **(C)** Similar analysis for Experiment 2 (diffuse attention condition; Materials and methods). Here, chance performance was at 0.33 proportion of correct trials, instead of 0.5 (see dashed horizontal lines in each panel). In both cases, peri-saccadic orientation identification performance was significantly higher in the upper rather than the lower visual field. Figures 2, 3 describe eye movement and visual stimulation controls that we analyzed in order to rule out other potential alternative explanations for different perceptual performance in the upper and lower visual fields. Also note that when computing the average peri-saccadic time course, we used a moving time window of 30 ms (Materials and methods), and we used window’s center when plotting. For example, the discrimination accuracy at time -7 ms included trials within the interval -22 to 8 ms (that is, the interval also included trials where the gabor grating was projected on screen while the eyes started moving towards the landing fixation point). We have added a visual representation of a single bin as a small grey box in panels B and C, centered at the time of -7 ms, to clearly showcase the temporal extent of the bin.

In the main experiments, each subject took part in two behavioral sessions, in non-consecutive days (day 1 and day 2). The experiment consisted of a gabor discrimination task, adapted from (Vallines and Greenlee, 2006). During the first session (day 1) each subject completed three training runs before a fourth experimental phase. Day 2 started directly with the experimental runs, without training runs. Each session lasted approximately 1.2 hours. The control fixation experiment included a single behavioral session of approximately 45 minutes.

On the first experimental day, the subjects first engaged in three training phases, each lasting approximately 4-5 minutes. For the first phase, the subjects were shown a fixation spot that jumped right or left by 12 deg, and they generated visually-guided saccades. We measured their baseline reaction times during this phase. For the second phase, the subjects maintained gaze fixation, and two image patches (like in Fig. 1A) were flashed for 1 frame (∼16.7 ms) either on the right or left side of the fixation spot (at a horizontal eccentricity of 6 deg). The patches were each at 3 deg above or below the horizontal meridian, and one of them was the target patch, while the other was the distractor patch. They both had a contrast of 40%. In Experiment 1, the target could have two orientations (horizontal or vertical), and in Experiment 2, it could have 3 orientations (horizontal, vertical, and oblique with direction +/-15 deg from the horizontal). Subjects practiced reporting the target orientation during fixation. Then, we moved to the third phase, in which we reduced the patch contrasts to 30% instead of 40%. We then started the main experiments.

Each experimental run consisted of 55 trials. Each subject took part in a variable number of experimental runs, ranging between 15 and 20 in two non-consecutive days. At the beginning of each run, a five point-calibration on the horizontal and vertical axes was performed. During each run, drift correction was applied every 7 trials. For each trial, subjects maintained central fixation and pressed the space bar to initiate a trial. After a variable interval between 750 ms and 1250 ms, the central fixation spot disappeared and a target fixation point was presented at 12 deg eccentricity, horizontally, randomly to the left or right with respect to central fixation. Subjects were asked to perform a saccade as quickly and as accurately as possible towards the target fixation point. At a variable interval from the requested saccade signal, we presented the target-distractor configuration (flash) on the same side as the requested saccade, for 1 frame or ∼16.7 ms (see Fig. 1A in Results) (Buonocore and Melcher, 2015; Fracasso et al., 2015; Fracasso and Melcher, 2016; Buonocore et al., 2017; Fabius et al., 2019; Fabius et al., 2020; Fabius et al., 2023).

The flash time interval was centered on the subject’s median saccadic reaction time estimated from the first training phase. We aimed at sampling behavioral performance around three main moments around the peri-saccadic interval: a) before saccade onset, presenting the target-distractor configuration 110 ms before the expected saccade onset time, as estimated from median saccadic reaction times; b) around saccade onset, presenting the target-distractor configuration at the expected saccade onset; and c) After saccade onset, presenting the target-distractor configuration 30ms after the expected saccade onset. This way, at the critical maximum saccadic suppression phase that we were interested in, we expected the stimuli to have appeared at approximately the vertical meridian (also see Figs. 2, 3 in Results). In Experiment 1, subjects reported one of two orientations as above, and in Experiment 2, they reported one of three orientations. Thus, Experiment 2 provided a more robust confirmation of Experiment 1’s results, since lucky guess rates were much less expected than in Experiment 1. Subjects were instructed to aim for accurate responses, not fast response times.

**Figure 2.**
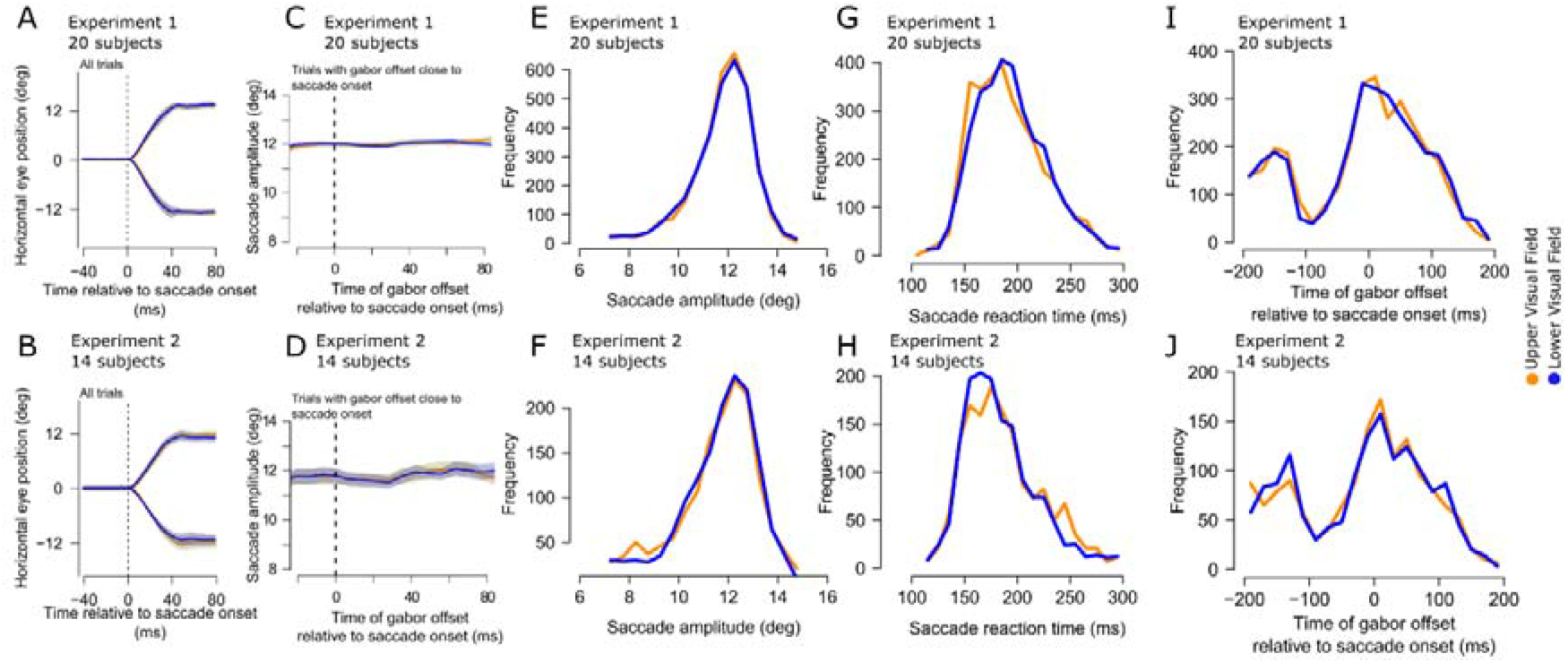
Similarity of eye movement metrics and timings between the upper and lower visual field target locations giving rise to differential peri-saccadic performance in **Fig. 1**. **(A, B)** Mean horizontal eye position traces for Experiment 1 **(A)** and Experiment 2 **(B)** separated by target location. Error bars denote two standard deviations. The saccades were similar whether the target appeared in the upper or lower visual field. **(C, D)** Saccade amplitude separated by target location along the peri-saccadic interval during which we observed the strongest differences in performance between the upper and lower visual fields. There was no systematic difference between the saccades for the different target locations. Saccadic amplitude **(E, F)** and reaction time **(G, H)** in each experiment were also similar for upper or lower visual field targets. **(I, J)** This implies that even the times of the gabor gratings relative to saccade onsets were matched between upper and lower visual field targets. Note that the dip in the histogram in each experiment is a known outcome of saccadic inhibition e.g. (Reingold and Stampe, 2002; Buonocore and McIntosh, 2008; Edelman and Xu, 2009; Bompas and Sumner, 2011), but it was, critically, no different between the two conditions. Thus, in all experiments, all eye movements were matched between upper and lower visual field target trials.

In each experiment, we had either a diffuse attention set of trials or a focused attention block of trials. In 50% of the experimental runs, the subjects were told that the target could either appear in the upper or lower visual field; their attention was thus diffuse, because they could not know a priori where the target was to be expected. In 25% of the runs, the subjects were told that the target will appear in the upper visual field with 97% probability; they could thus focus their attention on the upper visual field. And, in the final 25% of the runs, the subjects were told that the target will appear in the lower visual field with 97% probability, allowing them to focus their attention on the lower visual field stimulus. We randomly varied the order with which the diffuse and focused blocks of trials were run across individuals. That is, for some subjects, the diffuse block could start first followed by the two focused blocks, whereas for other subjects, one focused block could be finished first, then the diffuse block, and then the other focused block. Each subject was told which block they were running before they started their sessions.

On the control sessions, participants engaged with a training phase to familiarize themselves with the task. The subjects maintained gaze fixation, and two image patches (like in Fig. 1A of Results) were flashed for 1 frame (∼16.7 ms) along the vertical meridian, at either 3 deg or 7 deg in eccentricity. One of the patches was the target patch, while the other was the distractor patch. Target patches could be a 0.9 cpd gabor or a 6 cpd gabor. Gabor patches for the control experiment had a contrast of 15%. After the training phase, we collected real data for analysis with the same stimuli, allowing us to ask whether vertical meridian performance in our subjects was similar during gaze fixation (this control experiment) and peri-saccadically (Experiments 1 and 2). This is why the patches were presented vertically at 3 deg eccentricity in this control experiment (matching the retinotopic locations of flashes associated with maximal peri-saccadic suppression in Experiments 1 and 2). Each experimental run consisted of 45 trials. Each subject took part in a variable number of experimental runs, ranging between 7 and 9 in a single behavioral session.

### Human data analysis

Only trials in which a valid eye movement was executed entered the next stage of analysis. Valid eye movements had to be performed towards the landing fixation point and be between 7 deg and 15 deg in amplitude. Eye movement reaction time had to be between 100 ms and 300 ms (see Fig. 2 in Results), and with saccade duration shorter than 90 ms. For Experiment 1, 24% of trials were excluded based on these criteria, on average, across subjects. For Experiment 2, 32% of trials were excluded based on these criteria, on average, across subjects.

For perceptual reports, we computed the timing of the gabor offset relative to saccade onset by subtracting the time when the target-distractor configuration (flash) disappeared from the moment of saccade onset. According to this convention, negative values represent stimuli that were presented before the onset of the eye movement, while positive values represent stimuli that were (partially or in full) presented after saccade onset.

We also computed the distance traveled by the eyes while the target-distractor configuration (flash) was presented (‘Displacement of gabor on retina during flash’) by subtracting the eye position measured when the target-distractor configuration (flash) disappeared on screen from the eye position measured when the target-distractor configuration (flash) appeared on screen. This measure captured the distance traveled by the eyes over the target-distractor configuration, or the retinal slip of the flash, thus capturing potential saccade kinematic differences between experimental conditions that could account for discrimination performance during the peri-saccadic interval.

Finally, we computed the distance between the fovea and the target gabor when the target-distractor configuration disappeared from screen (‘Distance of gabor from fovea at flash offset’). This measure captured the distance between the fovea and the target gabor when the target-distractor configuration disappeared from screen, allowing us to assess potential differences in proximity of the fovea to the target gabor that could account for discrimination performance during the peri-saccadic interval. This measure, which we also complemented with careful analyses of subtle saccadic curvature, also motivated our placement of the gabor patches on the vertical meridian during the control experiment with gaze fixation.

Data were analyzed using the R software for statistical computing (Team). The data were analyzed with a Generalized Linear Mixed Model (GLMM) (Breslow and Clayton, 1993), based on the Generalized Linear Model framework proposed by McCullagh and Nelder (McCullagh and Nelder, 2019). Main effects and interaction between conditions for proportion of correct trials (binary outcome, 0-1) were tested using the logit function as link function (logistic regression model). Main effects and interaction between conditions for displacement of gabor on retina during flash and distance of gabor from fovea at flash offset were tested using the identity function (linear regression model). A subject numerical identifier was used as a random effect variable.

For some analyses, we also used a d’ measure of sensitivity, allowing us to rule out potential biases in the subjects’ reports. Because the d’ measure required including both hits and false alarms, we combined data from both focused and diffuse attention conditions in each experiment. This allowed us to stabilize the d’ measures, and it was fully justified because of the similarity of subjects’ performance under both attention conditions (see, for example, Fig. 5 in Results directly comparing focused and diffuse attention proportions of correct trials in each experiment).

To obtain time courses of orientation identification performance, for each participant, we used a moving time window of 30 ms, shifting its center by 7 ms at every iteration. Note that this means that for some time samples plotted in our Results figures, the 30 ms window spanned both a pre- and an intra-movement interval of measurement; we always clearly indicated these time samples in the figures and their legends. For every time window, we ran one GLMM for each dependent variable (proportion of correct trials, displacement of gabor on the retina during flash, and distance of gabor from fovea at flash offset, 3 models overall) and tested the main effect and interaction of the independent variables (gabor position and attentional state).

We used the Kolmogorov-Smirnov test to assess potential differences between distributions in the upper visual field and lower visual field experimental conditions for saccade reaction time, saccade gabor offset relative to saccade onset, and saccade amplitude.

In our experimental design, there was not a single-shot test for the joint intersection null (omnibus null) (García-Pérez, 2023). Our main null hypothesis did not consist of all comparisons between the upper and lower target positions being equal. Rather, we were particularly interested in orientation identification performance close to saccade onset, as well as while the eyes were moving towards the landing fixation point. Thus, we focused on the pairwise comparisons alone (García-Pérez, 2023). In all figures, we denoted asterisks for p-values less than 0.01 (Maier and Lakens, 2022). Otherwise, we also reported descriptive statistics and documented exact p-values when needed.

Finally, because our saccades were horizontal, vertical or horizontal gabor targets could be associated with different spatio-temporal energy as they traversed the retina (Castet et al., 2002; Schweitzer and Rolfs, 2020). Therefore, we had to rule out that higher peri-saccadic orientation identification performance in the upper visual field (see Results) was not trivially explained by a larger proportion of vertical peri-saccadic gabors in the upper visual field, which might be easier to visually discriminate. While this is unlikely due to our balanced experimental design, we checked this by calculating the proportion of vertical or horizontal gabors in every time window during which we analyzed orientation identification performance. There was no difference in the proportion of vertical or horizontal gabors between upper and lower visual field targets in the critical peri-saccadic intervals that we analyzed (see Results). This, along with our d’ analyses, ruled out potential biases in our subjects’ percepts due to different peri-saccadic gabor orientations relative to the saccade vector.

### Monkey neurophysiology

As we show in Results, our human experiments revealed a change of regime in orientation identification performance during peri-saccadic intervals (being better in the upper visual field when it is normally not so in the absence of saccades). Since the better orientation identification performance in the upper visual field is reminiscent of the SC’s preference for the upper visual field in its visual neural sensitivity (Hafed and Chen, 2016), we next asked how SC neurons behaved in peri-saccadic intervals. We analyzed the neuronal data presented in (Chen and Hafed, 2017) but from the perspective of visual field location. In that study, we documented saccadic suppression in the SC, but we did not explore effects of upper versus lower visual field locations. With this in mind, here we re-analyzed the same data from the perspective of visual field asymmetries, now asking whether saccadic suppression in the SC is different between the upper and lower visual fields. The behavioral and neurophysiological methods were described previously (Chen and Hafed, 2017).

Briefly, the monkeys fixated a small, central fixation spot. At some point during gaze fixation, a vertical gabor grating of different spatial frequencies (0.56, 1.11, 2.22, 4.44, and 11.11 cpd) and high contrast (100%) appeared within a recorded neuron’s RF and stayed there for a few hundred milliseconds. The monkeys were rewarded for simply maintaining fixation on the fixation spot until trial end. Because the stimulus stayed on for a prolonged period (unlike in the human experiments), we only analyzed trials in which the stimulus onset event happened after microsaccades. This interval is still an interval in which peri-saccadic suppression of the evoked visual burst still takes place (Hafed and Krauzlis, 2010; Chen and Hafed, 2017; Idrees et al., 2020a). Also, prior to running the main task, we mapped the RF’s using standard delayed and memory-guided saccade tasks. This allowed us to identify the RF hotspots and classify them as being in either the upper or lower visual field (Fig. 7A in Results). All microsaccades were also detected previously in the original study (Chen and Hafed, 2017). Here, we assessed their amplitude distributions across the upper and lower visual field sessions (Fig. 7B in Results), to ensure their similarity.

One main goal of the analysis was to investigate suppression of visual neural sensitivity around the time of microsaccades, and to determine if such modulation was different for neurons located in the upper or in lower visual field. To perform such analysis, for each neuron, we analyzed the neural activity following the stimulus onset in the mapping task, to determine the neuron’s RF hotspot location as the region of the visual field giving most activity. Once the hotspot was determined, upper visual field neurons were defined when the vertical component of the hotspot location was bigger than zero. All other neurons were labeled as lower visual field neurons. Then, we divided the data into two groups depending on whether saccades were executed or not during a critical interval around the stimulus presentation. In particular, no-saccade trials were defined as all the trials which did not have any saccades present between -100 to 100 ms around gabor onset. If a saccade was present in the time interval above, it was considered a saccade trial, and we assessed saccade time relative to stimulus onset time for evaluating time courses of neural suppression.

Spatial frequency tuning curves (i.e., responses for each given spatial frequency) were described previously (Hafed and Chen, 2016; Chen and Hafed, 2017), but in this, study we analyzed how saccades influenced these curves differently when the RF was either in the upper or lower visual field. To test the effect of saccadic suppression in the upper and lower visual fields, we computed a measure of “normalized firing rate”. First, we calculated, for each trial, the peak firing rate between 30 and 150 ms after stimulus onset. Then, for each neuron and spatial frequency condition, we averaged the peak firing rate in trials in which no saccades were detected. This value was then normalized by dividing the averages of each spatial frequency condition by the preferred spatial frequency response of that neuron, giving as a result the average tuning curve when no saccades were present. Similarly, for each neuron and spatial frequency, we averaged all the trials in which the gabor stimulus was presented 40 to 100 ms after saccade onset. The average peak firing rate at each spatial frequency condition was then normalized by the peak firing rate for the preferred spatial frequency response of the trials with no saccades. Doing so, values lower than one indicated suppression of neural activity because of saccade generation.

To summarize the time courses of saccadic suppression of SC visual bursts in the upper and lower visual fields (e.g. Fig. 9 in Results), we selected all the trials in which the gabor stimulus was presented between -50 to 140 ms relative to saccade onset. We then smoothed the data by applying a running average window of 50 ms on the normalized peak firing rate (relative to the baseline firing rate of for that spatial frequency) and by moving the average time window in steps of 10 ms. This analysis was performed only for the lower spatial frequency grating (0.56 cpd), which was the one used in the behavioral experiment reported above. To statistically test the difference between the upper and lower visual fields, we ran a series of two-sample independent t-tests at each bin of the two curves, and we set a threshold of 0.01 to determine whether a p-value was low enough to reject the null hypothesis.

## Results

### Peri-saccadic orientation identification performance is higher for upper visual field stimuli

We first asked whether human orientation identification performance around the time of saccades is different for upper or lower visual field peri-saccadic stimuli. In a first experiment (Experiment 1; diffuse attention condition), subjects generated approximately 12 deg horizontal saccades to the right or left of central fixation (Fig. 1A). At different times relative to saccade onset, a brief flash lasting approximately 16.7 ms was presented. The flash was centered horizontally at the midpoint between the initial fixation target location and the final desired saccade endpoint (that is, halfway along the intended saccade vector), and it consisted of two vertically-aligned image patches (each at 3 deg above or below the screen center). One patch was the target to be detected by the subjects, and it was either a horizontal or vertical gabor grating. The other patch was an irrelevant distractor without inherent orientation information (it was a superposition of two orthogonal gabors, with the total pattern tilted by 45 deg; Materials and methods). Across trials, the oriented patch was placed either above (upper visual field target location) or below (lower visual field target location) the horizontal meridian, and the other patch was at the vertically-symmetric position. There was also an equal likelihood of horizontal and vertical patches at each of the two locations. The subjects were instructed to report the orientation of the target flash (horizontal or vertical), and we assessed whether their performance differed as a function of target location.

Across 20 subjects, we found that peri-saccadic orientation identification performance was consistently better for upper visual field target locations when compared to lower visual field target locations. Specifically, Fig. 1B shows the time course of the proportion of correct trials in this experiment for targets flashed above (yellow) or below (blue) the horizontal meridian. During pre- and post-saccadic intervals long before or after the eye movements, performance was close to ceiling levels. However, in the critical peri-saccadic interval in which saccadic suppression was to be expected (Latour, 1962; Matin, 1974; Binda and Morrone, 2018), we found that the proportion of correct trials was significantly higher in the upper visual field than in the lower visual field (red asterisks; GLMM, main effect of target gabor location, p<0.01; see Materials and methods). Therefore, peri-saccadic orientation identification performance was significantly better in the upper visual field, unlike known lower visual field superiority of perceptual performance in the absence of saccades (Talgar and Carrasco, 2002; Montaser-Kouhsari and Carrasco, 2009; Barbot et al., 2021) (also see below for elaboration on this point with an explicit control experiment).

This result was also highly robust: we replicated the same observation in a second experiment (Experiment 2, diffuse attention condition), in which we increased task difficulty. Specifically, in this second experiment (Materials and methods), the target could have one of 3 different orientations, and we tested 14 subjects with it. The increased task difficulty allowed us to obtain a higher dynamic range of potential correctness results, minimizing ceiling and/or floor effects in the critical peri-saccadic interval. Once again, we found that orientation identification performance at the times near saccade onset (i.e. during peri-saccadic suppression) was consistently better for upper rather than lower visual field target locations (Fig. 1C, red asterisks; GLMM, main effect of target gabor location, p<0.01; see Materials and methods).

It is interesting to additionally note here that the difference in orientation identification performance between the upper and lower visual fields began to emerge even before saccade onset (Fig. 1B, C). While this observation is consistent with the fact that perceptual saccadic suppression does start before eye movement onsets in general (Diamond et al., 2000), we also highlight here that our temporal binning windows might have slightly amplified it (we used a moving window of 30 ms width in steps of 7 ms; see Materials and methods). Specifically, when plotting the average peri-saccadic time courses, we used the center of the moving time window as the sample being plotted, meaning that some time samples averaged both pre- and intra-movement intervals; an example sample centered on -7 ms is indicated by a shaded 30 ms bin in Fig. 1B, C to illustrate this point.

One potential caveat with the results of Fig. 1B, C so far could also have to do with the potential for the existence of subtle visual differences between the trials in which the target was presented either in the upper or lower visual field. We designed the flashes to always be symmetric around the horizontal meridian, minimizing visual differences between the upper and lower visual field trials. However, in control analyses, we also explicitly confirmed that the flashes appeared at similar times and retinotopic positions relative to the ongoing saccades, and that the saccades themselves were similar across the two conditions of upper versus lower visual field targets. Specifically, Fig. 2A, B shows the horizontal saccade trajectories in the two experiments for upper and lower visual field target positions. The trajectories were largely overlapping. The same conclusion holds true also for vertical saccade trajectories (not shown, but see Fig. 3 for equivalent evidence). Moreover, in Fig. 2C, D, we plotted the distributions of saccade amplitudes across time in the two conditions, with no differences in the saccade sizes between upper and lower visual field target trials, particularly along the peri-saccadic interval. Thus, the trials with upper or lower visual field targets had similar saccade trajectories, and therefore similar retinotopic visual stimulation by the targets. Similarly, saccadic amplitude (Fig. 2E, F), saccadic reaction times (Fig. 2G, H), and gabor offset times relative to saccade onset times (Fig. 2I, J) in the two experiments were always the same for upper and lower visual field targets. We confirmed this statistically: the distributions of saccadic amplitudes, saccadic reaction times, and gabor grating offset times relative to saccade onset times did not differ between trials with upper or lower visual field targets (Kolgomorov-Smirnov test, p>0.01). Therefore, the differences in peri-saccadic perceptual performance seen in Fig. 1 cannot be attributed to systematically different saccade parameters between upper and lower visual field target trials.

**Figure 3.**
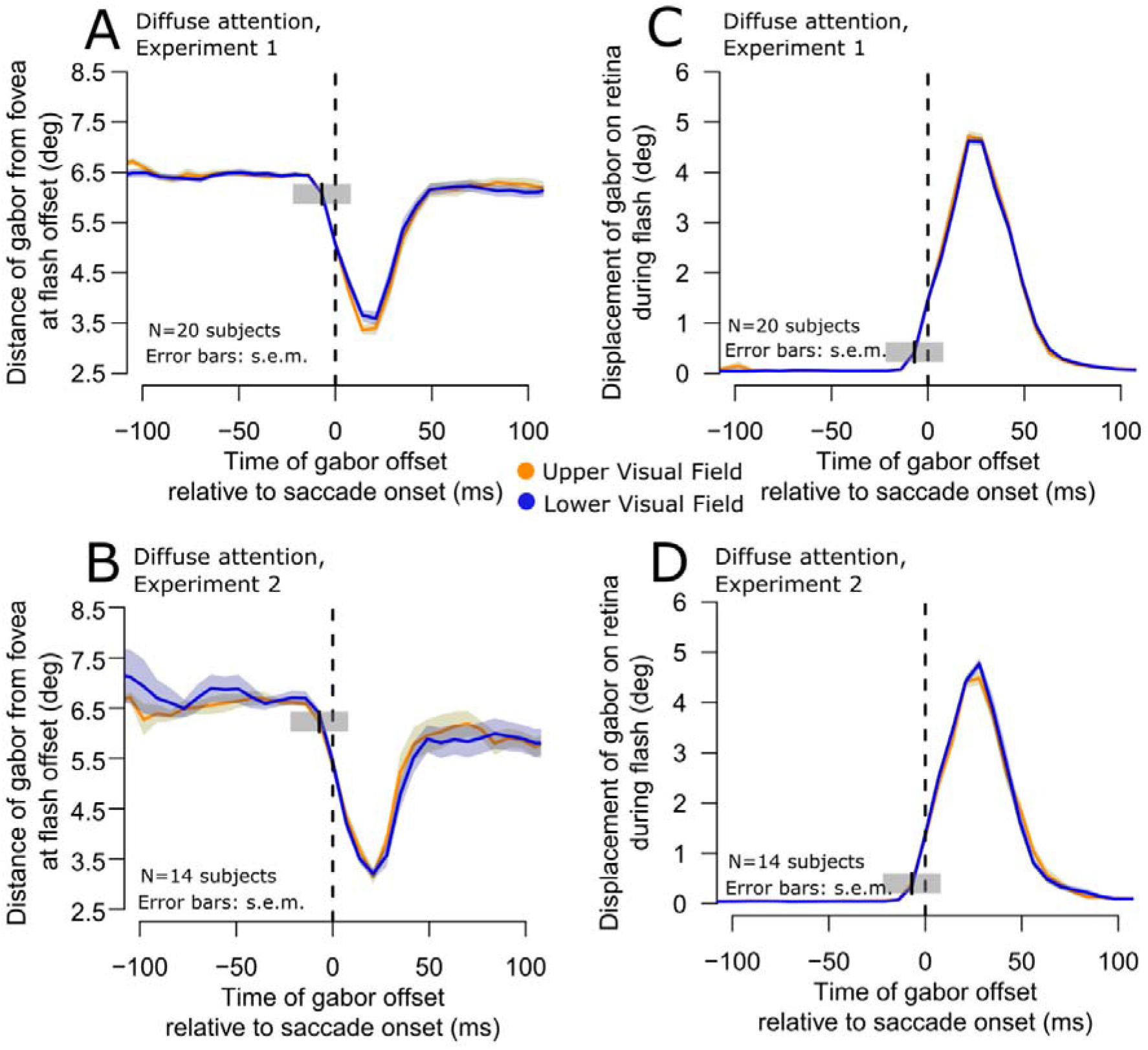
Similarity of retinal stimulation by the flashed gabor patches in the upper and lower visual field target trials. **(A, B)** In each experiment, we plotted the distance of the target gabor grating from the fovea as a function of time from saccade onset. During the saccade, the patches were closest to the fovea because the flash was always midpoint along the saccade path and timed to frequently occur peri-saccadically. However, and most critically, the distance to the fovea was not different for upper and lower visual field targets (compare yellow and blue curves in each panel). Therefore, the results of Fig. 1 were not due to a visual acuity benefit for upper visual field targets due to retinal eccentricity. Error bars denote s.e.m. **(C, D)** Similar analysis but for the retinal slip of the images during their onset (that is, the displacement of the gabor during its presentation). Because the eye was moving during a saccade, the grating slipped in position on the retina. However, once again, such retinal slip was the same for upper (yellow) and lower (blue) visual field targets in both experiments. Note that the gray shaded regions with center at time -7 ms are similar to those shown in Fig. 1, and they illustrate the extent of our binning window for one example sample point. They also explain why the retinal slip may have appeared to start increasing even before saccade onset (this was only a consequence of our temporal binning windows and not because of erroneous saccade onset detection).

We also considered whether potential saccadic curvature might have differed sufficiently between the two conditions to influence the results of Fig. 1. That is, it could be argued that the retinotopic position of the flash might have been systematically closer to the fovea for upper versus lower visual field target flashes (perhaps due to saccadic curvature), which would have conferred a slight orientation identification advantage for the upper visual field targets. However, this was again not the case. In Fig. 3A, B, we plotted the distance of the gabor grating from the fovea at the time of its offset in the two conditions (upper versus lower visual field target locations), and in the two experiments. There was clear overlap in this distance between the two target locations. Moreover, since the flash sometimes happened during the eye movements themselves, we also plotted the retinal slip of the flash in Fig. 3C, D. Again, such slip was similar whether the target flash was in the upper or lower visual fields, and this was the case in both experiments. Similar results were obtained in the focused attention (Materials and methods) trials as well.

Furthermore, saccades that depart slightly from a horizontal trajectory, due to subtle curvature, might have an impact on the foveal availability of stimuli presented in the periphery in either the upper or lower visual field. For this reason, we further analyzed our data separately for leftward only saccade trials or rightward only saccade trials (even though the curvatures of the two saccade directions were not statistically significantly different from each other), and we observed virtually indistinguishable results between saccade directions.

Therefore, the retinal conditions of the flashes were similar for upper and lower visual field targets, meaning that the results of Fig. 1 were not trivially explained by systematically different retinotopic stimulation between conditions.

Finally, gabors orthogonal to the executed saccade can potentially be harder to resolve than parallel gabors (Castet et al., 2002; Schweitzer and Rolfs, 2020). Hence, a difference between the proportion of horizontal and vertical gabors in the lower and upper visual field target conditions could potentially also bias our results. While this is unlikely given our balanced experimental design (Materials and methods), to control for this possibility, we additionally analyzed the proportion of horizontal gabors along the peri-saccadic interval and across the tested experimental conditions; we did this using the same time course techniques adopted for orientation identification performance (Fig. 1), saccadic amplitude (Fig. 2C, D), distance of gabor from the fovea, and displacement of gabor on retina (Fig. 3). In Experiment 1, the proportion of trials in which a horizontal gabor was presented hovered around 50% in the interval from -100 ms to around 90 ms, as could be expected from a balanced two-alternative forced choice task, with no discernible difference between trials presented in the upper or lower visual field. In Experiment 2, the proportion of trials in which a horizontal gabor was presented hovered around 33%, compatible with a three-alternative forced choice task. Also in this case, no systematic difference between trials presented in the upper of lower visual field could be observed. Thus, our results in Fig. 1 cannot be trivially accounted for by a systematic occurrence of more targets that are easier to discriminate in the upper versus lower visual field.

To summarize, our results so far demonstrate that peri-saccadic orientation identification performance is significantly higher in the upper rather than lower visual field. These observations complement prior work by Knöll et al. (Knöll et al., 2011), who have documented the topography of saccadic suppression along the horizontal meridian. These authors found that peri-saccadic suppression occurred in a retinotopic frame of reference, with a divisive reduction of sensitivity that was constant across the retinal eccentricity dimension. To our knowledge, peri-saccadic orientation identification performance in the upper and lower visual fields, around the time of a horizontal eye movement, has not been reported previously.

### Valid prior knowledge of upper or lower visual field target location does not alter the result

Perhaps the strongest evidence that better upper visual field peri-saccadic orientation identification performance was a robust phenomenon emerged when we gave our subjects, within contiguous blocks of trials, valid prior knowledge about the upcoming target location. Specifically, in approximately one quarter of all trials in each experiment (Materials and methods), the subjects were explicitly told that the current block of trials had primarily only upper visual field targets (with 97% probability). Similarly, in another one quarter of the trials, the subjects were informed that the current block of trials had primarily lower visual field target locations (with 97% probability). We called these blocked trials the focused attention trials. In both cases, orientation identification performance in the peri-saccadic interval was still higher in the upper visual field than in the lower visual field. This result is shown in Fig. 4. That is, even when the subjects fully knew in advance that a target was going to appear in the lower visual field, their peri-saccadic orientation identification performance to such a target was still lower than the orientation identification performance for targets in the upper visual field. Note also that eye movement control analyses in the focused attention conditions (Figs. 2, 3) again ruled out any eye movement or retinal stimulation explanations of the results. Thus, even valid advance knowledge of target position did not eliminate the observation of stronger peri-saccadic orientation identification performance in the upper visual field.

**Figure 4.**
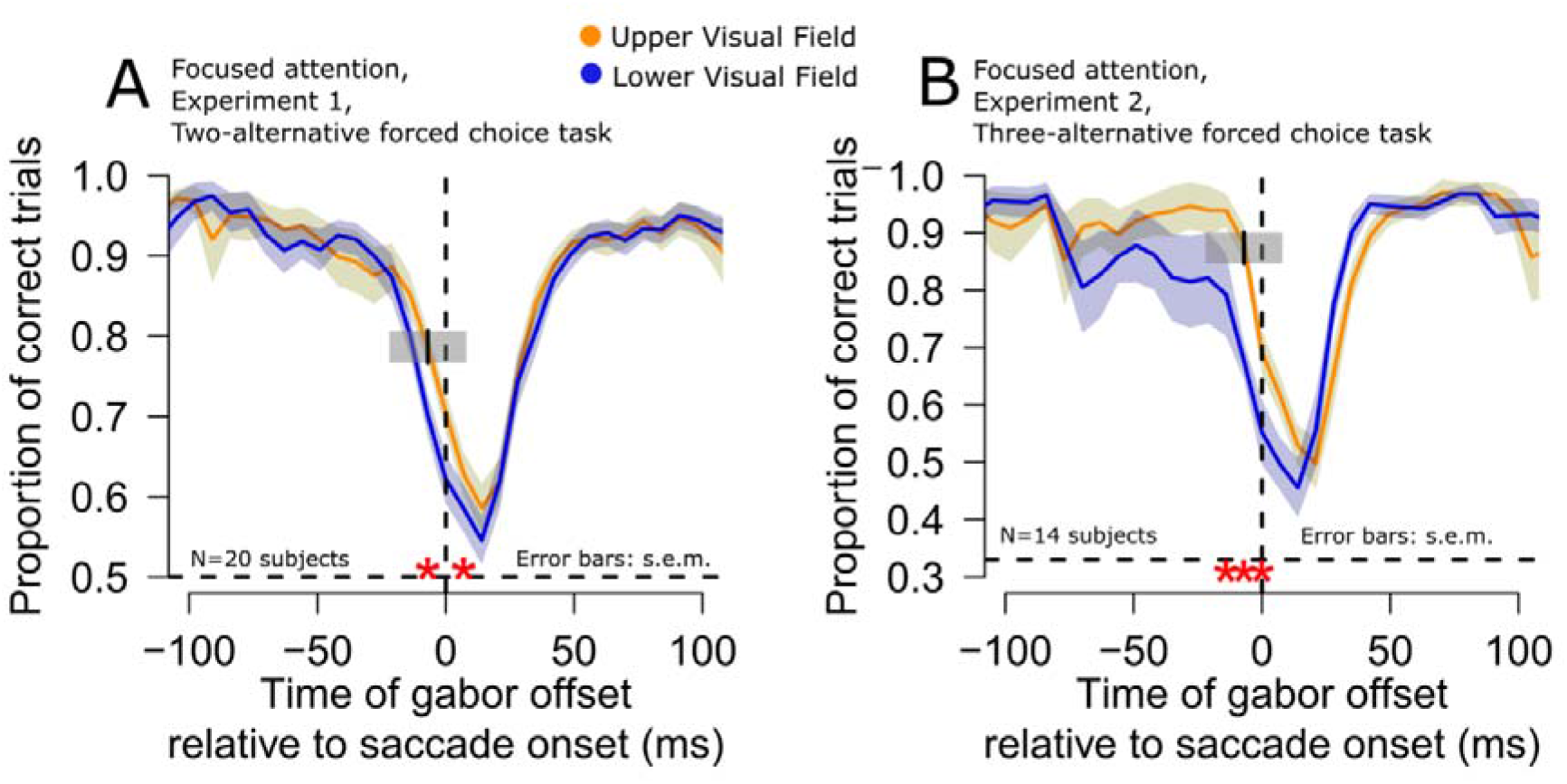
Persistence of the upper visual field peri-saccadic orientation identification advantage even with full advance prior knowledge of target location. **(A, B)** In both experiments, peri-saccadic upper visual field orientation identification performance was still better than for the lower visual field (red asterisks) even with valid prior knowledge of target location (Materials and methods). All other conventions are similar to those in Fig. 1.

Of course, the results of Fig. 4 were not entirely only a negative result with respect to the blocking manipulation of target position. For example, when we compared orientation identification performance long before saccade onset (−200 to - 70 ms from saccade onset) in the diffuse and focused attention conditions, both experiments were suggestive of a perceptual benefit when prior knowledge about target location was provided. For example, in Experiment 1, the subjects exhibited 88% average correct rates with prior knowledge of target location (focused attention trials) when compared to 86% average correct rates without prior knowledge (p=0.055, paired t-test). In Experiment 2, the average correct rates were 91% and 88% in the diffuse and focused attention trials, respectively (p=0.017, paired t-test). Therefore, the lack of influence of advanced prior knowledge on peri-saccadic orientation identification performance alluded to above (Fig. 4) was primarily restricted to the peri-saccadic interval.

It is important to note here that while it is not possible to perform a direct comparison between performance in Experiment 1 and Experiment 2 (one was a two-alternative and the other was a three-alternative forced choice paradigm), we did collect both diffuse and focused attention trials within each of these experiments (Materials and methods). Therefore, we also had an opportunity to compare peri-saccadic focused and diffuse attention effects within each experiment. For this reason, we plotted orientation identification performance within the +/-20 ms interval around saccade onset, summarizing this comparison (Fig. 5). In both experiments, peri-saccadic orientation identification performance was significantly higher in the upper rather than the lower visual field, irrespective of attention condition. Moreover, within each experiment, the effect size for the focused and diffuse attention conditions was similar, and no interaction between target location and attention condition was observed. In fact, statistical analysis in each experiment (including both diffuse versus focused trial comparisons) revealed that there was neither a main effect of attentional instruction nor an interaction effect between gabor grating position and attentional instruction (Figs. 4, 5A, B, red asterisks; GLMM, main effect of target gabor location, p<0.01; see Materials and methods). Similar conclusions were also made when we calculated d’ based on all trials within each experiment in the intervals shown in Fig. 5 (Materials and methods).

We also note here that m-AFC tasks force observers to give a response on every trial, and authentic correct responses might be mixed up with lucky guesses arising in cases in which the observer could not gather or use sensory information to give an informed response. For future measurements it could be interesting to allow observers to give “don’t know” responses instead of producing a guess on trials in which they have no basis for responding, and explicitly model those guess responses, while fitting the authentic judgements (García-Pérez and Alcalá-Quintana, 2019; Reynolds et al., 2021). In the current measurements, we were fully cognizant of the need to decrease noise in the data. That is why we ran our second experiment, employing a 3-AFC task and limiting the issues associated with guessing by the subjects; the larger number of choices in this task variant leads to a lower chance performance level, which can be reached faster (i.e. within fewer trials), compared to 2-AFC designs (Jäkel and Wichmann, 2006).

### Orientation identification performance is not higher in the upper visual field during fixation

In a control experiment, we next explicitly tested whether our results could arise simply during fixation as well, in which case there would be nothing special about the peri-saccadic results reported above. We required participants to keep fixating the center of the screen while gabors were presented in the upper or lower visual field (on the vertical meridian). Thus, retinotopically, the gabors were at a similar location to that in the peri-saccadic interval at maximal perceptual suppression (Figs. 2, 3). This provided a better comparison than testing orientation identification performance in the main experiments long before saccade onset, since the gabors were not on the retinotopic vertical meridian long before the saccades in the main experiments (Fig. 1A).

In this control experiment, the gabors could also either have a high (6 cpd) or low (0.9 cpd) spatial frequency, with the latter being the same spatial frequency that was used in Experiments 1 and 2. We also presented the gabors either close to the fovea (3 deg, mimicking the minimal distance between the center of the gabor and the fovea while the eyes were moving towards the saccade landing position; Fig. 3) or far from the fovea (7 deg). The results indicated the expected benefit for the lower visual field during fixation (Carrasco et al., 2001; Kristjansson and Sigurdardottir, 2008), specifically for stimuli with high spatial frequency (p < 0.01; Fig. 6); this was decidedly opposite our peri-saccadic observations. For the low spatial frequency, there was neither a benefit nor a cost associated with the lower visual field, consistent with (Kristjansson and Sigurdardottir, 2008). Importantly, we did not observe the peri-saccadic upper visual field benefit that we observed with saccades in any of the tests that we conducted involving gaze fixation. Thus, the peri-saccadic results of Figs. 1-5 above were indeed qualitatively different from those expected during gaze fixation.

**Figure 5.**
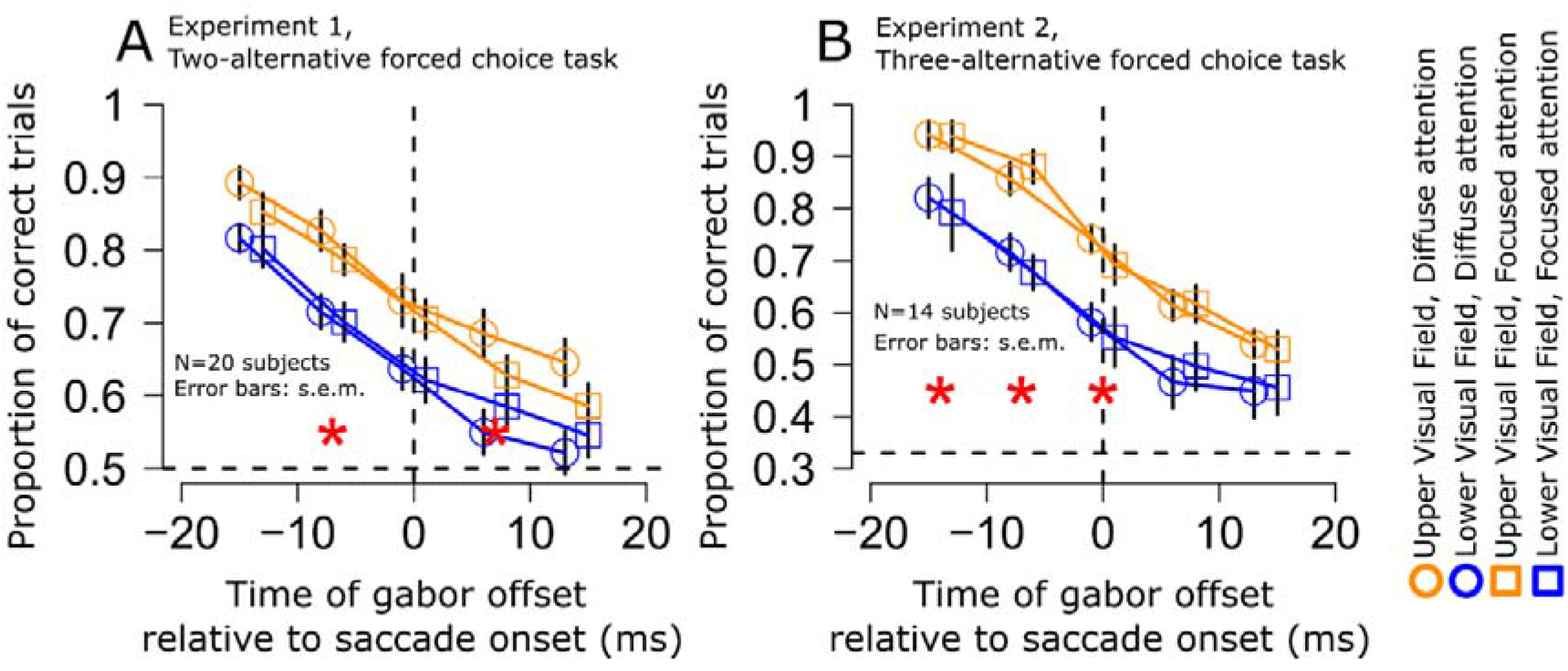
Similarity of peri-saccadic orientation identification performance between the focused and diffuse attention conditions in each experiment. **(A)** Time course of orientation identification performance relative to saccade onset for targets in the upper (yellow) or lower (blue) visual field in Experiment 1 (diffuse attention condition: circles; focused attention condition: squares; see Materials and methods). Red asterisks indicate significant differences between upper and lower targets (GLMM, main effect of target gabor grating location, p<0.01). **(B)** Similar analysis for Experiment 2. Here, chance performance was at 0.33 proportion of correct trials, instead of 0.5 (see dashed horizontal line). In both cases, peri-saccadic perceptual performance was significantly higher in the upper rather than the lower visual field. It is important to note that within each experiment, the effect size for the focused and diffuse attention conditions was similar, and no interaction between target location and attention condition was observed.

**Figure 6.**
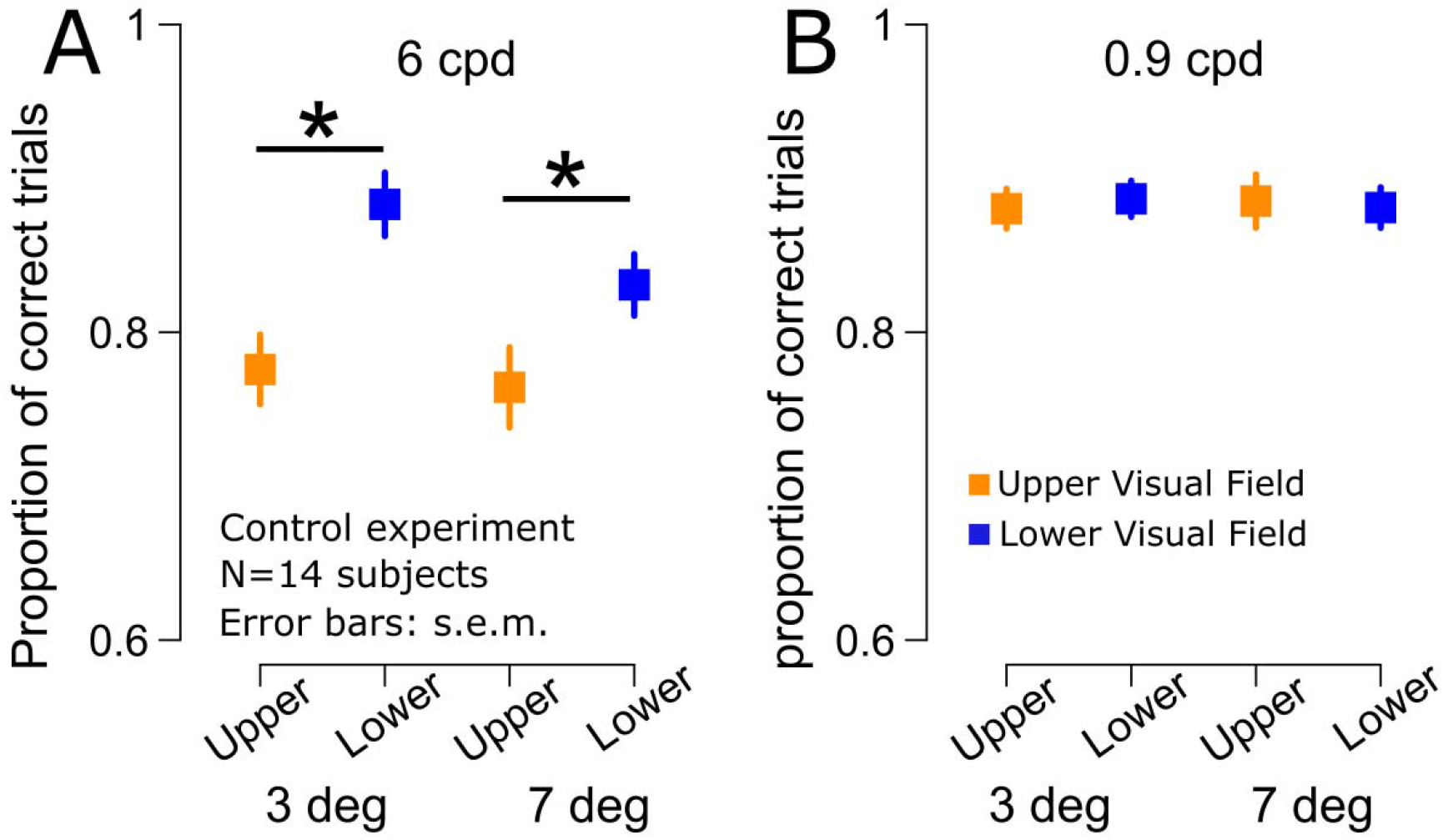
Lack of upper visual field advantage in orientation identification performance during fixation, in the absence of saccades. **(A)** Discrimination performance at fixation for high spatial frequency gabors (6 cpd) presented in the upper and lower visual fields (along the vertical meridian) at 3 deg or 7 deg eccentricity. We observed a significant benefit for gabors presented in the lower visual field at both 3 and 7 deg (asterisks indicate p < 0.01). **(B)** Discrimination performance at fixation for low spatial frequency gabors (0.9 cpd) presented in the upper and lower visual fields (along the vertical meridian) at 3 deg or 7 deg eccentricity. There was neither an upper nor lower visual field advantage, but our peri-saccadic results (with 0.9 cpd) showed a clear upper visual field advantage. The results in this figure mimic closely those reported in prior publications during fixation (Carrasco et al., 2001; Kristjansson and Sigurdardottir, 2008), and they confirm that our observed peri-saccadic asymmetries in performance (e.g. Figs. 1, 4, 5) are indeed qualitatively different from those expected during fixation.

### Stimulus-induced superior colliculus visual responses to peri-saccadic stimuli are also higher in the upper visual field

Our results so far suggest that peri-saccadic orientation identification performance in humans is better in the upper visual field, a result that is qualitatively different from how performance of visual tasks normally manifests during maintained gaze fixation (Talgar and Carrasco, 2002; Montaser-Kouhsari and Carrasco, 2009; Barbot et al., 2021). This implies that peri-saccadic orientation identification performance may be dominated by pathways other than the classic visual cortical systems exhibiting anisotropies favoring the lower visual field (Benson et al., 2021; Kupers et al., 2022). Interestingly, unlike the visual cortex, the SC does preferentially process upper, rather than lower, visual field stimuli during fixation (Hafed and Chen, 2016). However, it is still not known whether this still holds peri-saccadically. Therefore, we next checked whether stimulus-induced SC firing rates to peri-saccadic stimuli were still stronger in the upper visual field. We speculate that this might provide a putative mechanism consistent with our interpretation of the perceptual results above (e.g. Fig. 1).

To explore this hypothesis, we conducted a second, separate study in which we analyzed the visual responses of 115 SC neurons collected from neurophysiological recordings performed on awake monkeys. We knew from prior work that these neurons exhibited robust saccadic suppression for stimuli appearing immediately in the wake of microsaccades (Chen and Hafed, 2017). We chose this particular dataset (Materials and methods) to carefully analyze for visual field asymmetries because of two primary reasons. First, microsaccades are an effective means to study saccadic suppression in the SC (Hafed and Krauzlis, 2010; Chen and Hafed, 2017) because microsaccades are genuine saccades (Hafed et al., 2009; Hafed, 2011), and because they have the advantage of not moving visual response fields (RF’s) too much due to their small size. Therefore, presenting stimuli to the RF’s with and without the rapid eye movements (to assess suppression relative to baseline) is experimentally simple with microsaccades. Second, in this data set, we used stimuli presented directly in the post-movement interval after the microsaccades (Chen and Hafed, 2017), allowing us to avoid (as much as possible) the visual effects of retinal image displacements during the movements themselves.

We first assessed that the recorded neurons were similarly distributed across the upper and lower visual fields. Figure 7A shows the RF hotspot directions in deg, relative to the horizontal cardinal axis, for all of the recorded neurons. Negative numbers indicate neurons representing the lower visual field, and positive numbers indicate neurons with RF hotspots above the horizontal meridian. From the figure, it can be seen that the two populations of neurons were equally sampled across the upper and lower visual fields. Similarly, in Fig. 7B, we plotted the amplitudes of microsaccades occurring near stimulus onset (and thus associated with peri-saccadic suppression), which were similar in the sessions in which we recorded neurons with either upper or lower visual field RF’s. Therefore, the eye movement characteristics were similar regardless of whether we recorded upper or lower visual field SC neurons.

**Figure 7.**
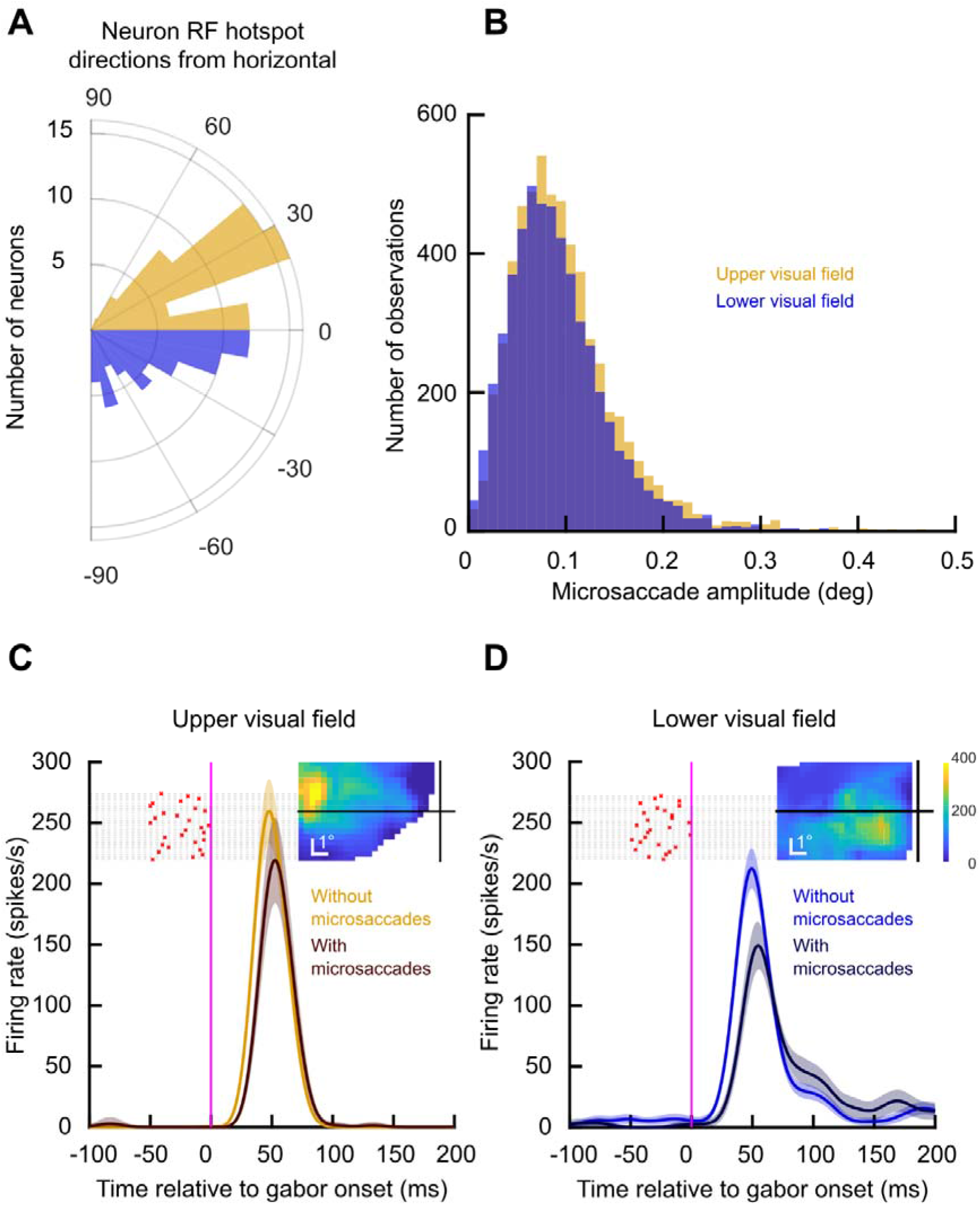
Higher upper visual field peri-saccadic sensitivity in SC neurons. **(A)** Distribution of RF hotspot locations from our recorded population, expressed as a direction from the horizontal meridian. Approximately half of the neurons had RF hotspots in the upper visual field (yellow), and the rest had hotspots in the lower visual field. **(B)** We assessed peri-saccadic suppression by evaluating visual sensitivity for stimuli appearing near the time of microsaccades (Hafed and Krauzlis, 2010; Chen and Hafed, 2017). Here, we characterized the microsaccade amplitudes for the two groups of sessions that we compared (in which we recorded either upper or lower visual field SC neurons). The eye movement amplitudes were matched across the two groups. **(C)** Example upper visual field SC neuron responding to the onset of a low spatial frequency gabor grating (0.56 cpd). The saturated yellow curve shows the neuron’s visual response in the absence of nearby microsaccades (Materials and methods), and the darker curve shows the same neuron’s visual response when the stimulus appeared immediately in the wake of microsaccades (individual microsaccade onset times are shown as a trial raster of red crosses in the background of the figure). The inset shows the RF location of this neuron, indicating that it preferentially represented a part of the upper visual field. Error bars denote 95% confidence intervals. **(D)** Similar to **C** but for a neuron preferring the lower visual field (see RF map in the inset). Not only did the neuron have lower baseline visual firing rate (saturated blue curve) (Hafed and Chen, 2016), but its suppressed visual response (darker curve) was also more strongly reduced than in the neuron in **C**. Thus, SC visual burst strength was still higher in the upper visual field for peri-saccadic stimuli.

When we then inspected the neurons’ visual responses themselves, we observed consistently higher SC firing rates in the upper visual field neurons than in the lower visual field neurons for peri-saccadic stimuli. Consider, for example, the pair of neurons shown in Fig. 7C, D. In Fig. 7C, the neuron had an upper visual field RF (the RF map is shown in the inset). Its visual response to a low spatial frequency grating of 0.56 cpd was mildly suppressed when the grating appeared immediately after microsaccades. Specifically, the yellow curve shows the neuron’s average firing rate in the absence of microsaccades near stimulus onset (Materials and methods), and the darker curve shows the average firing rate when the grating appeared immediately after microsaccades (individual microsaccade times across different trials from this condition are shown as red crosses in the figure). The neuron’s response was suppressed in association with microsaccades, as expected but such suppressed response was still robust and peaking above 200 spikes/s. On the other hand, the neuron in Fig. 7D represented a lower visual field location (its RF map is shown in the inset). Not only was its baseline visual response (in the absence of nearby microsaccades) weaker than the baseline response of the neuron in Fig. 7C (Hafed and Chen, 2016), but its peri-saccadically suppressed response (dark curve) was also more strongly affected by the eye movements. In other words, the neuron experienced stronger saccadic suppression than the neuron in the upper visual field, consistent with our perceptual results above. Thus, if anything, the spatial anisotropy in the SC in terms of upper versus lower visual field neural sensitivity (Hafed and Chen, 2016) was amplified even more during peri-saccadic intervals.

We confirmed this by isolating a measure of saccadic suppression and confirming that it was stronger for lower rather than upper visual field SC neurons. Across the population, we normalized each neuron’s activity by its strongest no-microsaccade visual response to any of the five different spatial frequencies that we tested (Chen and Hafed, 2017); that is, we picked the spatial frequency that evoked the strongest peak response, and we normalized all trials’ firing rate measurements by this value (Materials and methods). We then normalized each neuron’s peri-saccadically suppressed visual response using the very same normalization factor, and we averaged across neurons. For the neurons preferring the upper visual field (Fig. 8A), the population generally preferred low spatial frequencies (Chen et al., 2018) in its baseline no-microsaccade activity (yellow curve; error bars denote 95% confidence intervals). However, the tuning curves were broader than in the lower visual field neurons (Fig. 8B, blue curve). For example, the upper visual field neurons were more sensitive to 4.44 cpd gratings than the lower visual field neurons, consistent with prior observations (Hafed and Chen, 2016). Most importantly for the current study, for the peri-saccadically suppressed visual bursts (dark curves in Fig. 8), similar observations persisted. That is, the upper visual field neurons had broader tuning curves than the lower visual field neurons in the peri-saccadic interval, and they were suppressed less than the lower visual field neurons at the low spatial frequencies. For example, at the lowest spatial frequency (0.56 cpd), there was weaker saccadic suppression in the upper visual field neurons (Fig. 8A) than in the lower visual field neurons (Fig. 8B); this is evidenced by the larger difference between the blue and dark blue curves in Fig. 8B than between the yellow and dark yellow curves in Fig. 8A (p=0.038, two-sample t-test).

**Figure 8.**
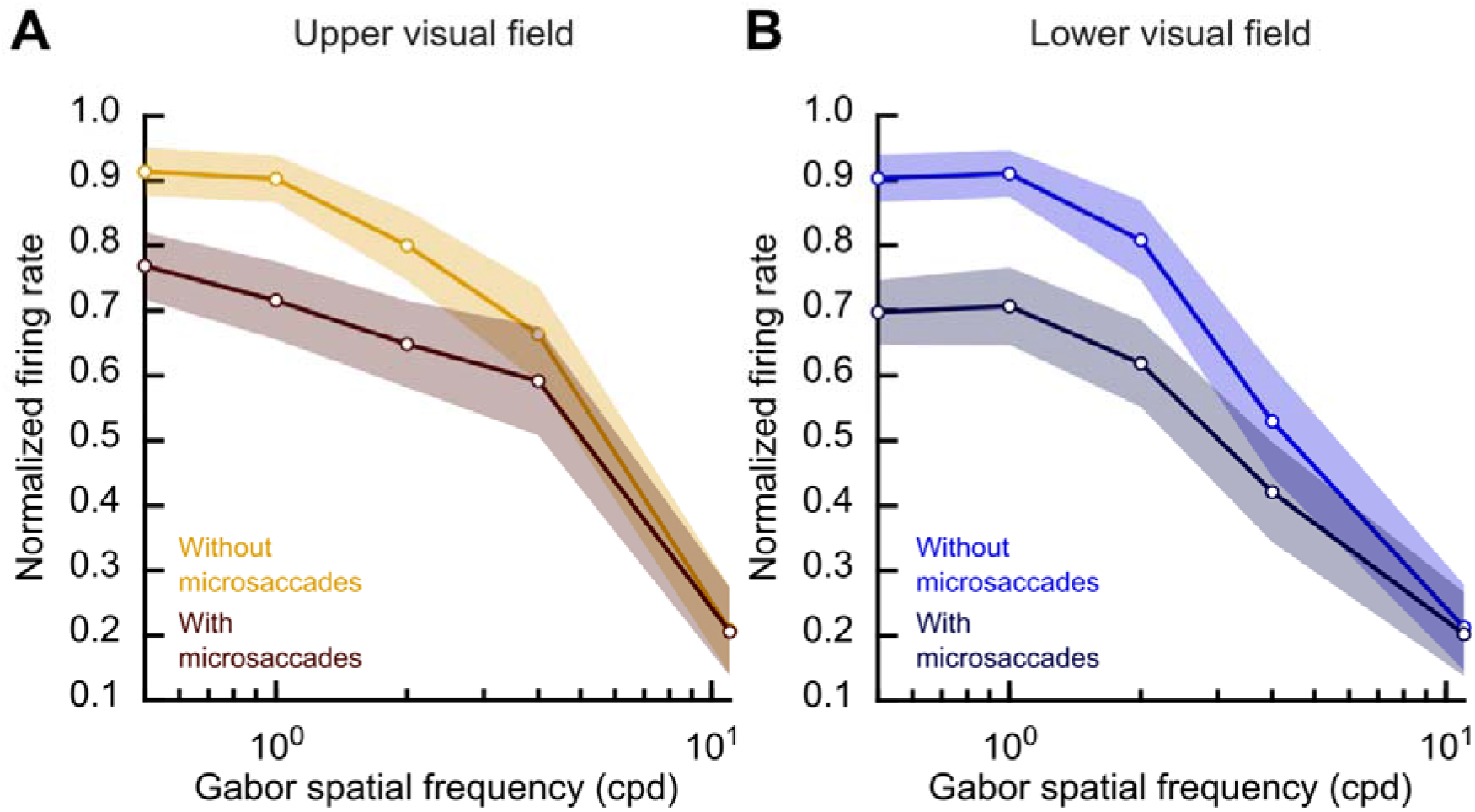
Broader peri-saccadic SC population tuning curves in the upper visual field. **(A)** Average population tuning curve of the upper visual field neurons without eye movements (saturated yellow) and peri-saccadically (dark). Error bars denote 95% confidence intervals. In both curves, we normalized each neuron’s firing rate to the peak visual response for the preferred spatial frequency (Materials and methods). Lower spatial frequencies experienced more suppression for peri-saccadic stimuli than higher spatial frequencies, as expected (Chen and Hafed, 2017). **(B)** Same analysis but for the lower visual field neurons. In baseline (saturated blue), the neurons were more low-pass in nature than the upper visual field neurons in **A** (Hafed and Chen, 2016). For example, the tuning curves dropped sharply at 4.44 cpd when compared to the neurons in **A**. This difference persisted for the peri-saccadic tuning functions (that is, there was stronger saccadic suppression in the darker curve when compared to **A**); also see Fig. 7.

At higher spatial frequencies, the saccadic suppression effect was expectedly weakened overall (Chen and Hafed, 2017), but this weakening again happened more so for the upper visual field neurons than for the lower visual field neurons (for example, the difference between the curves at 2.22 and 4.44 cpd was smaller in the upper visual field, panel A, than in the lower visual field, panel B). Coupled with the fact that the neurons were themselves more sensitive in the upper visual field in the no-microsaccade trials (Hafed and Chen, 2016) (e.g. Fig. 7), this suggests that there was a consistently higher firing rate in the SC visual bursts in the upper visual field for peri-saccadic stimuli.

We also confirmed the above interpretations by plotting the neural peri-saccadic suppression time course profiles, like we did for the human experiments above. We found consistently higher relative firing rates in the upper visual field neurons than in the lower visual field neurons, as can be seen in Fig. 9 for the case of 0.56 cpd grating stimuli. To obtain this figure, we calculated the normalized firing rate for each trial in which the gabor grating appeared in the interval from -50 ms to 140 ms relative to movement onset (see Materials and methods). We then plotted the mean normalized firing rate at each time bin for neurons in the upper (yellow) versus lower (blue) visual fields. Values lower than one indicated a reduction in firing rate, which took place for both upper and lower visual field neurons (indicating peri-saccadic suppression). Most critically, and consistent with Figs. 7, 8, the peak suppression was stronger by about 10% for the neurons in the lower visual field (blue) compared to the neurons in the upper visual field (yellow; p<0.01). Similar trends were observed for higher spatial frequencies, but they got progressively weaker and weaker as expected from Fig. 8 and ref. (Chen and Hafed, 2017). We conclude that SC peri-saccadic visual neural sensitivity is consistently higher in the upper visual field.

**Figure 9.**
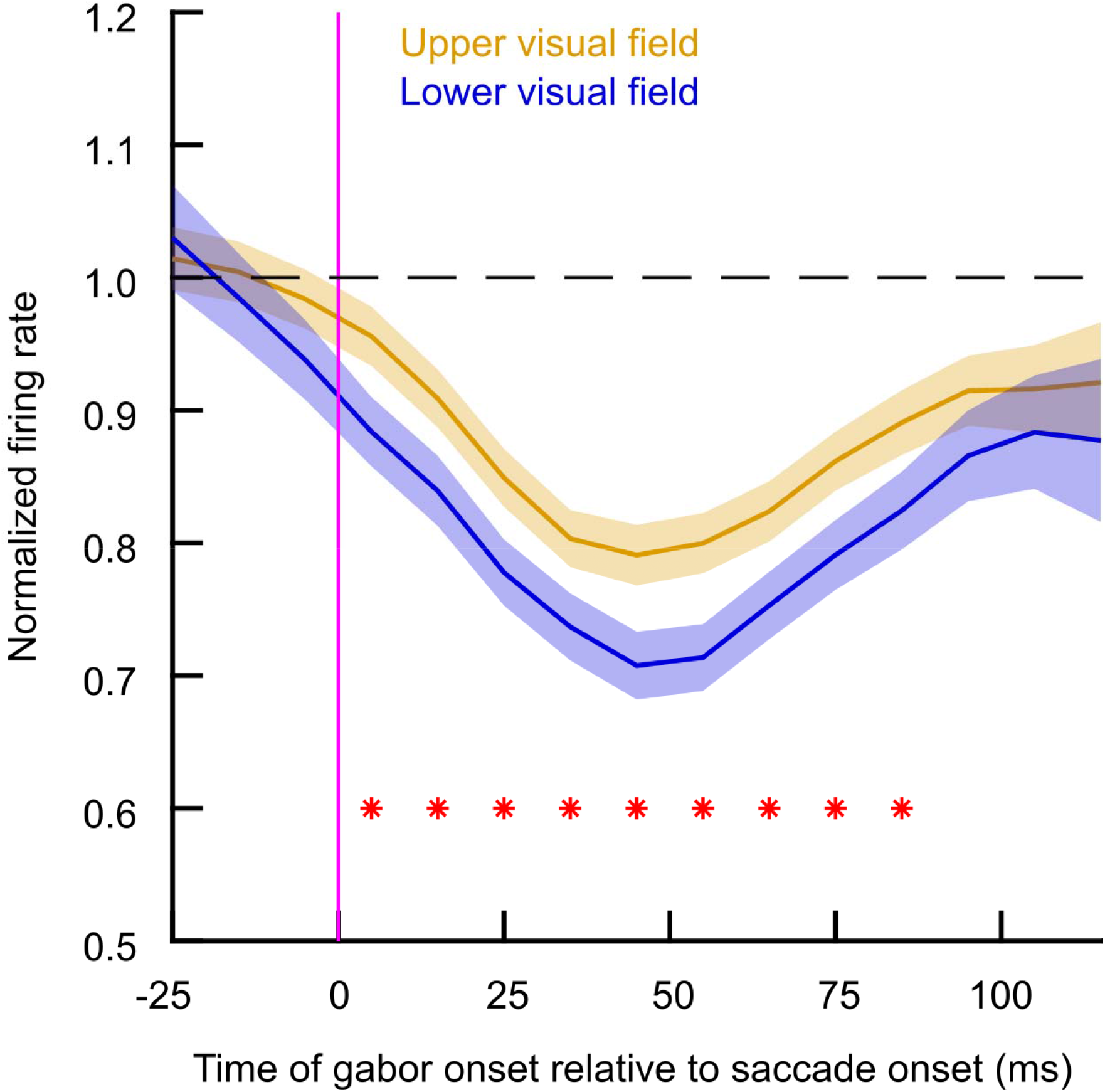
Milder suppression in upper visual field SC neurons in peri-saccadic times. The figure shows the time course of normalized visual neural responses in the SC for low spatial frequencies (Materials and methods). Upper visual field neurons (yellow) experienced milder saccadic suppression, and were therefore more sensitive, than lower visual field neurons (blue). Such an effect was temporally locked to the saccades, as in Fig. 1B, C. Thus, coupled with overall higher visual sensitivity in upper visual field SC neurons, these results suggest that during peri-saccadic intervals, the anisotropy between upper and lower visual field SC sensitivity is even larger than in the absence of eye movements.

## Discussion

In this study we observed paradoxically better orientation identification performance for peri-saccadic visual stimuli presented in the upper compared to the lower visual field (Figs. 1, 3). That is, at the time of strong peri-saccadic perceptual suppression (Diamond et al., 2000; Idrees et al., 2020b), perceptual performance violated a well-known observation that lower vertical meridian vision could be significantly better than upper vertical meridian vision (Himmelberg et al., 2020; Barbot et al., 2021) (also see Fig. 6). Moreover, peri-saccadic visual responses in the monkey SC were stronger in the upper visual field, and therefore qualitatively correlated with the human orientation identification performance. We speculate that this observation might provide a starting point for an account of our behavioral results, and it can motivate future neurophysiological studies directly testing both behavioral and neural modulations in the very same subjects.

Our SC results are particularly interesting. Visual neural sensitivity in the SC is relevant for perceptual performance, as supported by numerous studies showcasing a relationship between SC visual response properties and saccadic reaction times (Boehnke and Munoz, 2008; Marino et al., 2012; Marino et al., 2015; Hafed and Chen, 2016; Chen et al., 2018; Malevich et al., 2022). For example, it has consistently been shown how neuronal sensitivity (Boehnke and Munoz, 2008; Marino et al., 2012; Marino et al., 2015; Hafed and Chen, 2016; Chen et al., 2018) and neuronal latency (Chen et al., 2018) can successfully predict visual detection performance as reported by reflexive orienting responses. Thus, our observations could suggest that peri-saccadic vision may prioritize readiness to orient in general, which could be useful for taking corrective actions after the executed saccade. Of course, it is important to note here that more direct measures of orientation identification performance in monkeys would be the next experimental step when exploring neurophysiological mechanisms.

Activity in the SC has also been strongly linked to overt and covert attentional shifts, probed behaviorally with landolt-C visual acuity stimuli (Ignashchenkova et al., 2004). Moreover, the SC is relevant for motion, form, and spatial frequency discrimination (Sprague et al., 1970; Lovejoy and Krauzlis, 2010; Chen et al., 2018). And, complex visuo-motor behavior in the SC has been critically linked to the ability to discriminate between real and self-induced movements (Robinson and Wurtz, 1976). Most importantly, a causal inactivation study has specifically demonstrated a perceptual deficit after inactivating the SC (Lovejoy and Krauzlis, 2017). And increasing evidence demonstrates that SC neurons are even sensitive to high-level stimuli like faces (Bogadhi and Hafed, 2022); also see (Hafed et al., 2023). Thus, it was important for us to document SC results here.

Having said that, different visual sources could contribute to the effects that we observed. The target stimuli that we adopted were all relatively small, confined to the upper or lower visual field, and briefly presented around the onset of the eye movement. Such stimuli are often mislocalized along directions related to the eye movement vector (Honda, 1989; Lappe et al., 2000) and also visually suppressed. Interestingly both suppression and mislocalization could reflect visual mechanisms. For example, suppression could arise due to retinal-image motion during the saccade and the high-contrast images preceding and following the saccade (Wurtz, 2008; Garcia-Perez and Peli, 2011). Similarly, there exists evidence for mislocalization asymmetries, with stronger mislocalization reported for upward saccades than for other saccade directions; such asymmetries could be accounted for, at least theoretically, by how visual images are translated peri-saccadically on the warped representation of the visual image in the SC (Grujic et al., 2018). Thus, there exist differences in mislocalization patterns for the upper and lower visual fields that could in principle correlate with the finding we observed in the current manuscript.

From a visual suppression perspective, other authors (Garcia-Perez and Peli, 2011) have compared contrast sensitivity in upper and lower visual fields during saccades and fixation, and they have not found systematic differences between upper and lower visual fields across individuals. Thus, low-level contrast processing does not appear to account for our findings.

When considering active vision, the existence of spatial anisotropies in neural circuits and behavior is intriguing, in general. In particular, not only is visual performance generally better in the lower visual field (Himmelberg et al., 2020; Barbot et al., 2021), but attentional performance is as well (He et al., 1996; Rubin et al., 1996). Moreover, cortical visual areas may have anisotropies that are in line with such behavioral anisotropies favoring the lower visual field (Benson et al., 2021; Kupers et al., 2022). However, for the oculomotor system, opposite anisotropies exist. The SC strongly favors upper visual field visual stimuli (Hafed and Chen, 2016). Moreover, saccades are faster (Schlykowa et al., 1996; Zhou and King, 2002; Hafed and Chen, 2016; Hafed and Goffart, 2020) and more accurate in the upper visual field (Hafed and Chen, 2016), the latter likely reflecting significantly smaller movement fields in the SC upper visual field representation. However, if that is indeed the case, how does vision operate during peri-saccadic intervals? We found that it behaves more like the oculomotor anisotropy, as in being better in the upper visual field, than the visual cortical anisotropy. This dichotomy is interesting to consider from a broader perspective, especially when discussing more general questions regarding the role of the SC in cognition in general. For example, increasing evidence suggests that the SC may be a controller of visual attentional modulations in the cortex (Lovejoy and Krauzlis, 2010; Hafed et al., 2011, 2013; Lovejoy and Krauzlis, 2017; Krauzlis et al., 2018; Bogadhi et al., 2019; Bogadhi et al., 2021). However, if this is the case, then how might one reconcile the opposite anisotropies that the SC and visual cortices exhibit?

One possibility might be that the pattern of feedback that the SC provides to the cortex is combined to serve either attention or perceptual performance at strategic times. For example, it may be the case that larger visual RF’s in the lower visual field representation of the SC aid multiple smaller RF’s in the cortex to be functionally bound together during directed covert attention to a given location (that is, without saccades). This could jointly modulate the normally separate cortical RF’s. Thus, the opposite anisotropy between the SC and visual cortex may actually be functionally useful during gaze fixation. In the case of peri-saccadic vision, the opposite anisotropy may be useful in an additional manner: to favor detecting far, extra-personal stimuli (e.g. aerial threats) exactly at the time in which perception may be most compromised by saccadic suppression. This can aid in quick orienting or evasive responses. Thus, it may be favorable to have better peri-saccadic vision in the upper visual field, like in the SC, than in the lower visual field, like in the cortex. This, in turn, might mean that the gain of feedback from the SC to the cortex, which may be useful for saccadic suppression (Berman et al., 2017; Baumann et al., 2022), is higher for lower visual field locations than upper visual field locations (that is, causing stronger saccadic suppression).

We find this idea useful, and plausible, in placing our results in the context of other recent observations related to active vision. For example, we recently found that SC saccade-related bursts are stronger in the lower visual field, not the upper visual field (Hafed and Chen, 2016; Zhang et al., 2022). Interestingly, saccade kinematics were not different for upper and lower visual field saccades, suggesting that the SC motor bursts do not necessarily dictate movement kinematics (Zhang et al., 2022). Instead, we think that they may modulate the gain of feedback to the cortex, perfectly supporting our observations of stronger saccadic suppression in the lower visual field. Indeed, there is evidence that feedback projections from the SC to the frontal cortex may target inhibitory neurons (Sommer and Wurtz, 2004; Shin and Sommer, 2012), and inactivation of the SC during saccades renders saccade-related frontal cortical bursts stronger rather than weaker (Berman et al., 2009). All of this evidence suggests that there may be asymmetric gain feedback to the cortex from the SC, which causes stronger saccadic suppression in the lower visual field. One prediction of the above idea, therefore, is that we should also observe stronger neuronal peri-saccadic visual sensitivity in the upper visual field in cortical visual areas, not just in the SC, but this idea remains to be tested.

Another interesting insight from our SC results is that within the SC itself, the visual anisotropy between the upper and lower visual fields is magnified peri-saccadically. That is, not only are neurons less sensitive to visual stimuli presented in the lower visual field (Fig. 7), but they also experience stronger saccadic suppression in the peri-saccadic intervals (Figs. 8, 9). Therefore, the already strong disparity in neuronal visual sensitivity between the upper and lower visual fields in the SC (Hafed and Chen, 2016) is rendered even stronger peri-saccadically.

Finally, it is interesting to consider that even full prior knowledge of target location (Figs. 4, 5) did not necessarily alter our observations in the perceptual experiments. This suggests that fundamental mechanisms governing peri-saccadic vision operate under practically all conditions, irrespective of attention. This might have a useful ecological purpose, as mentioned above. At the time during which vision is most compromised by saccades, it might be most useful to utilize whatever remaining residual visual abilities, under all behavioral contexts, to detect extra-personal stimuli (which primarily reside in the upper visual field) and rapidly react to them.

## Acknowledgements

The authors would like to thank Isla Macvicar, Noor Musaed N Alsedairi, Oisean Burnett, Rebecca Taylor and Rosanne Timmerman. A.F. was supported by a grant from the Biotechnology and Biology Research Council (BBSRC, grant number: BB/S006605/1) and the Bial Foundation (Bial Foundation Grants Programme; Grant id: A-29315, number: 203/2020, grant edition: G-15516). A. B. and Z. M. H. were funded by the Deutsche Forschungsgemeinschaft (DFG) under projects HA6749/2-1 (FOR 1847; project A6) and BU4031/1-1.

